# Plant signals anticipate the induction of the type III secretion system in *Pseudomonas syringae* pv. *actinidiae* facilitating efficient temperature-dependent effector translocation

**DOI:** 10.1101/2022.06.01.494460

**Authors:** Maria Rita Puttilli, Davide Danzi, Cristiana Correia, Jessica Brandi, Daniela Cecconi, Marcello Manfredi, Emilio Marengo, Conceição Santos, Francesco Spinelli, Annalisa Polverari, Elodie Vandelle

**Author notes:** These authors contributed equally to this work (Maria Rita Puttilli, Davide Danzi).

## Abstract

Disease resistance in plants depends on a molecular dialogue with microbes that involves many known chemical effectors, but the time course of the interaction and the influence of the environment are largely unknown. The outcome of host–pathogen interactions is thought to reflect the offensive and defensive capabilities of both players. When plants interact with *Pseudomonas syringae*, several well-characterized virulence factors contribute to early bacterial pathogenicity, including the type III secretion system (T3SS), which must be activated by signals from the plant and environment to allow the secretion of virulence effectors. The manner in which these signals regulate T3SS activity is still unclear. Here, we strengthen the paradigm of the plant–pathogen molecular dialogue by addressing overlooked details concerning the timing of interactions, specifically the role of plant signals and temperature on the regulation of bacterial virulence during the first few hours of the interaction. Whole-genome expression profiling after 1 h revealed that the perception of plant signals from kiwifruit or tomato extracts anticipates T3SS expression in *P. syringae* pv. *actinidiae* compared to apoplast-like conditions, facilitating more efficient effector transport *in planta*, as revealed by the induction of a temperature-dependent hypersensitive response in the non-host plant *Arabidopsis thaliana* Col-0. Our results show that, in the arms race between plants and bacteria, the temperature-dependent timing of bacterial virulence versus the induction of plant defenses is probably one of the fundamental parameters governing the outcome of the interaction.

**Abstract importance:** Plant diseases– their occurrence and severity– result from the impact of three factors: the host, the pathogen, and the environmental conditions, interconnected in the disease triangle. Time was further included as a fourth factor accounting for plant disease, leading to a more realistic three-dimensional disease pyramid to represent the evolution of disease over time. However, this representation still considers time only as a parameter determining when and to which extent a disease will occur, at a scale from days to months. Here, we show that time is a factor
regulating the arms race between plants and pathogens, at a scale from minutes to hours, and strictly depends on environmental factors. Thus, besides the arms possessed by pathogens and plants per se, the opportunity and the timing of arms mobilization should make the difference to determine the outcome of an interaction and thus the occurrence of plant disease.

## Introduction

The course and outcome of host–pathogen interactions are thought to be determined mainly by the ability of each player to adapt existing or newly acquired genetic resources that provide enhanced offensive or defensive capabilities (1). The genes and effector molecules involved in such interactions have been investigated in the hope of developing new strategies for disease control in plants, but this has yielded few practical improvements. The comparative analysis of bacterial genomes has identified many genes involved in plant–microbe interactions, which provide insight into bacterial host range and virulence. However, further investigation is required to understand the dynamics of gene expression during interactions and the functions of effector proteins in both partners. In the interaction between a host plant and *Pseudomonas syringae*, several virulence factors contribute to early bacterial pathogenicity. For successful infection, *P. syringae* relies on a large repertoire of virulence effectors, which mainly act to suppress plant immunity and construct a favorable niche for bacterial growth within the apoplast (2). Such effectors are translocated into host cells via the type III secretion system (T3SS) (3), which requires an activating signal to achieve competence. Comparative microarray analysis based on publicly available genome data revealed that the *hrp/hrc* gene cluster (encoding the T3SS) is strongly induced in biovar 3 of the kiwifruit bacterial canker pathogen *Pseudomonas syringae* pv. *actinidiae* (Psa3), but only weakly induced in biovar 1 (Psa1) and unaffected in biovar 2 (Psa2), revealing differences in transcriptional responsiveness to apoplast-like conditions among the three biovars, particularly in the context of T3SS activation (4). Moreover, a reporter gene construct combining the promoter of the *hrpA1* gene (encoding one of the main components of the T3SS pilus) with the gene for green fluorescent protein (GFP) as an indicator of Psa virulence induction, confirmed that one or more signals present in kiwifruit extracts boost *hrpA1* promoter activity in Psa3 (4). Although Psa1 induces weaker T3SS-related gene expression in minimal medium compared to Psa3, it responds similarly to kiwifruit extracts, suggesting the presence of common sensors in the two biovars. In contrast, the *hrpA1* promoter remained inactive in Psa2 under all the conditions tested (4).Psa is therefore an ideal model for studies involving the induction of bacterial T3SS-related genes during interactions with plants, because the three biovars respond differently to minimal growth medium and/or kiwifruit extracts.

Although the T3SS activation process is poorly understood, there is compelling evidence that activation occurs following the perception of target cells (5,6). Accordingly, the *hrp* gene cluster in *P. syringae* is controlled by plant signals and multiple physiological and environmental factors, specifically pH, osmotic potential, and catabolite repression. All *hrp/hrc* genes are induced *in vitro* under conditions that simulate the leaf apoplast environment, widely described as “minimal medium” (7,8). Moreover, the *hrp* genes of the plant pathogen *Ralstonia solanacearum* are under the genetic control of *hrpB*, a regulatory gene whose expression is induced when bacteria are co-cultivated with plant cell suspensions, due to the recognition of unidentified non-diffusible signals in the plant cell wall (9). Despite the evidence described above, transcriptomic analysis with bacteria grown under different conditions does not always show the induction of T3SS-related genes *in planta* or in the presence of plant extracts *in vitro* (10,11). In some cases, the absence of *hrp* gene induction may result from the cultivation of bacteria in minimal medium, which already represents optimal conditions for the induction of T3SS-related genes. Alternatively, the inability to detect *hrp* gene induction *in planta* may indicate that induction is transient and subsides 48–72 h after inoculation. Indeed, large-scale transcriptome profiling experiments have mostly represented samples prepared several hours after infection or cultivated for several hours in the presence of plant extracts, thus obscuring the very early stages of the interaction.

To address the lack of data covering the early stages of plant–microbe interactions, we profiled the transcriptome of Psa3 grown in apoplast-like medium, which revealed the induction of T3SS-related gene expression after only 1 h in the presence of kiwifruit or tomato leaf extract. This suggests that interaction with the host plant anticipates the T3SS-related gene expression already induced by apoplast-like conditions, as confirmed by an increase in the abundance of secreted proteins. Moreover, given that phytopathogenic bacteria must overcome plant defenses that suppress T3SS activity, the induction of T3SS-related gene expression by plant extracts suggests that Psa enhances the activity of T3SS *in planta*, as revealed by the temperature-dependent hypersensitive response observed in the non-host plant *Arabidopsis thaliana* Col-0.

## Results

### Kiwifruit and tomato extracts enhance gene expression in Psa3 after only 1 h

The treatment of Psa3 with kiwifruit or tomato extracts led to the modulation of several genes after only 1 h compared with growth in minimal medium. We detected 73 DEGs (72 upregulated and 1 downregulated) in the presence of kiwifruit extract and 186 DEGs (127 upregulated and 59 downregulated) in the presence of tomato extract (**Supplementary Datasets S1, S2**). Among the upregulated genes, 33 were common to both treatments, whereas 39 were specifically induced by the kiwifruit extract and 94 by the tomato extract (**Fig. 1A**). The genes showing a statistically significant induction in response to the extracts were analyzed using STRING to identify clusters of associated proteins, revealing multiple functional clusters involved in a variety of pathophysiological processes. Among the genes induced by kiwifruit extract, the most obvious STRING cluster contained 25 proteins associated with the T3SS and type III effectors (**Fig. 1A**, red and purple circles, respectively, left panel). Among the genes induced by tomato extract, STRING analysis revealed proteins involved in translation and components of the Sec system, representing ~50% of the upregulated genes (**Fig. 1A**, red circle, right panel). The other two functionally related groups featured proteins associated with sulfur metabolism (yellow circle, right panel) and galactose metabolism (green circle, right panel). Finally, the 33 genes upregulated by both extracts included several encoding type III effectors and cognate chaperones as well as the type III helper protein HopP1 (**Supplementary Dataset S3**).

**Fig. 1.**
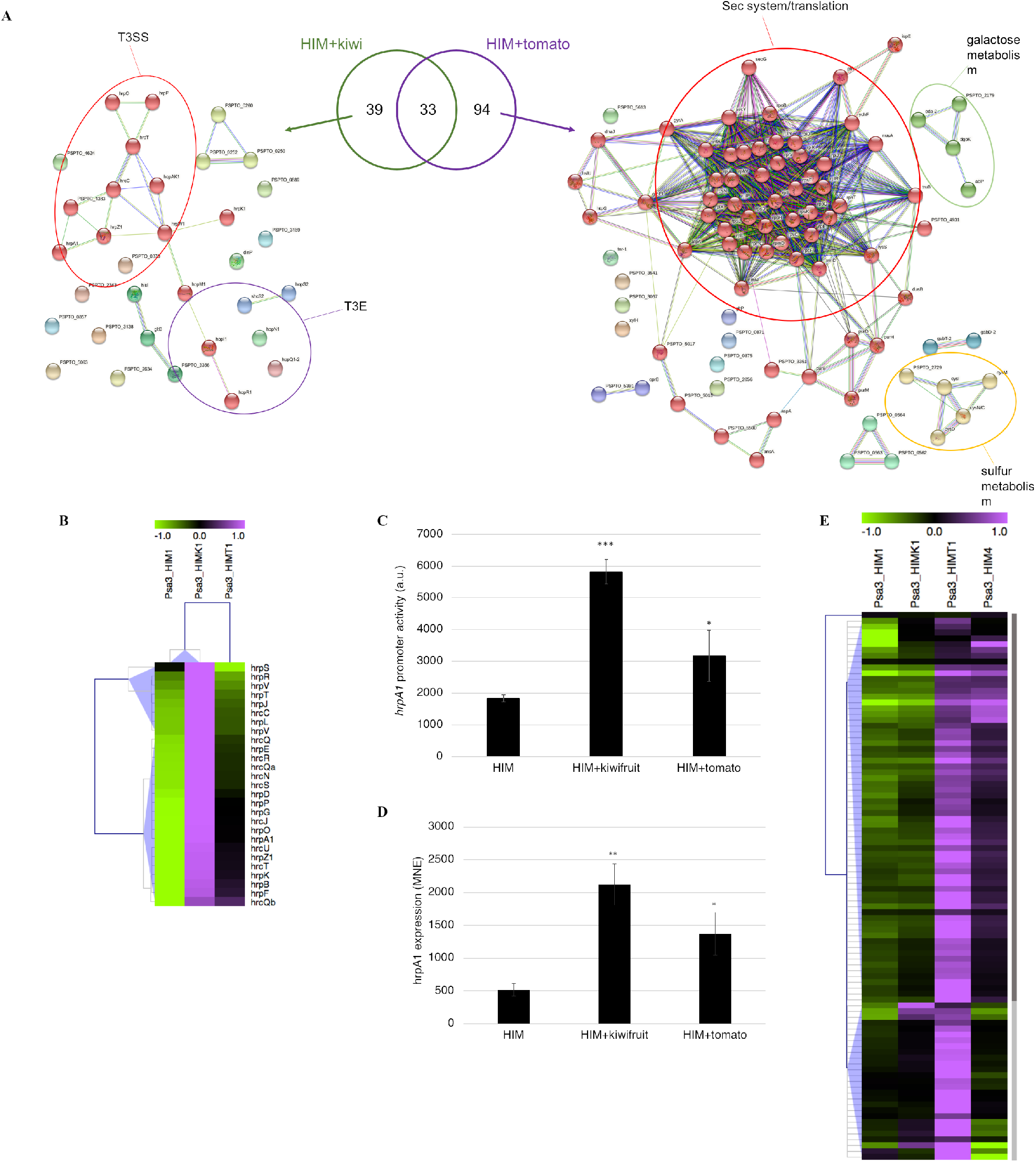
Kiwifruit and tomato extracts upregulate virulence-related genes after only 1 h in *P. syringae* pv. *actinidiae* (Psa) grown under apoplast-like conditions. (A) Common and specific genes upregulated in Psa3 by kiwifruit (left panel) and tomato (right panel) extracts and their functional relationship analyzed using STRING. The main categories (T3SS = type III secretion system, T3E = type III effectors) are highlighted with colored circles. The Venn diagram, generated using Draw Venn Diagram, shows common and unique upregulated genes for the two treatments. (B) Hierarchical clustering of the absolute expression levels of Psa3 *hrp*/*hrc* genes after incubation for 1 h in minimal (*hrp*-inducing) medium (HIM1), or HIM supplemented with kiwifruit (HIMK1) or tomato (HIMT1) extract. (C) Activity of the *hrpA1* promoter in a Psa3 strain carrying the *phrpA1::gfp* reporter system after incubation for 1 h in HIM or HIM supplemented, as indicated, with kiwifruit or tomato extract. (D) Expression of *hrpA1* in Psa3 after incubation for 4 h in HIM or HIM supplemented, as indicated, with kiwifruit or tomato extract. Gene expression was analyzed by realtime qPCR using *rpoD* as a housekeeping gene for normalization, and is expressed as mean normalized expression (MNE). (E) Hierarchical clustering of the absolute expression level of genes upregulated by tomato extract in Psa3 grown in HIM for 1 h (HIM1) or 4 h (HIM4), or HIM supplemented with kiwifruit (HIMK1) or tomato (HIMT1) extract. In (B) and (E), clusters were generated using MeV with normalized fluorescence values, and blue triangles indicate the different clusters. In (C) and (D), the asterisks indicate a statistically significant differences between treated and untreated samples according to a Student’s t-test (**p < 0.05, * p < 0.1).

We focused on the *hrp/hrc* genes, the major category of Psa3 genes induced by kiwifruit extract. Although every gene in the *hrp/hrc* cluster was not differentially expressed, the transcripts were nevertheless always more abundant in the presence of plant extracts compared to minimal medium (HIMK1 *vs* HIM1 and HIMT1 *vs* HIM1; **Fig. 1B**). The significance of this induction was confirmed by analyzing *hrpA1* promoter activity in a bacterial strain carrying the construct p*hrpA1*::GFP, as well as *hrpA1* expression by real-time qPCR. These experiments confirmed that *hrpA1* is upregulated in the presence of each extract, although to a different extent (**Fig. 1C,D**). The genes induced only by the tomato extract (Psa3_HIMT1) were generally upregulated more strongly than those induced only by the kiwifruit extract (Psa3_HIMK1), but the transcripts in the latter group were still more abundant compared to cells incubated for 1 h in minimal medium (Psa3_HIM1) (**Fig. 1D**). Most of these transcripts also became more abundant when the incubation period was extended to 4 h (Psa3_HIM4). These results indicate that the extracts share common molecules that can induce bacterial signaling pathways including the T3SS, but the quantity of the shared molecules varies. Interestingly, the genes upregulated by kiwifruit and tomato extracts were also induced in Pto DC3000 grown *in planta* for 6 h (19) (**Supplementary Fig. S1**), suggesting the genes play a functional role during the first phase of interaction, as expected for the T3SS, supporting the use of plant extracts as a model to study the influence of plant signals on bacterial behavior.

### Kiwifruit extract anticipates the Psa3 response and triggers the faster induction of bacterial virulence

To determine whether the genes induced by kiwifruit extract after 1 h are specifically associated with the presence of plant signals, we compared these genes to those induced after 4 h in apoplast-like medium with (HIMK4) or without (HIM4) host plant extract. The resulting Venn diagram shows that 68% (49/72) of the genes induced by kiwifruit extract after 1 h were still induced after 4 h (HIMK4) compared to 1 h in minimal medium (HIM1) (**Fig. 2A**), and 43 of them were also induced after 4 h in the absence of kiwifruit extract (HIM4 *vs* HIM1; **Supplementary Dataset S4**). Moreover, compared to 1 h in apoplast-like medium (HIM1), 121 genes were upregulated in the presence and absence of kiwifruit extract, representing ~70% (HIMK4 *vs* HIM1) and ~92% (HIM4 *vs* HIM1) of the induced genes, respectively. Hierarchical clustering of all DEGs identified under these conditions revealed three major clusters with different profiles (**Fig. 2B**). In cluster I, transcript levels were modulated specifically in the presence of kiwifruit extract after 1 h. In cluster II, transcript levels increased after 4 h regardless of the presence or absence of kiwifruit extract. Finally, the larger cluster III contained genes that were induced in the presence of kiwifruit extract after 1 h, followed by a further increase in expression after 4 h regardless of the presence or absence of the kiwifruit extract. Interestingly, although Gene Ontology enrichment revealed no significant categories in clusters I or II, the genes in cluster III were significantly enriched for protein transport/secretion functions involving the T3SS (**Fig. 2B**, right panel), suggesting that kiwifruit signals do not induce T3SS (and related effectors) but rather anticipate the upregulation that would normally occur after 4 h in minimal medium. In line with this hypothesis, no DEGs were identified in Psa3 following incubation for 4 h in the presence or absence of kiwifruit extract, confirming that the genes were still induced under apoplast-like conditions, but only after a delay.

**Fig. 2.**
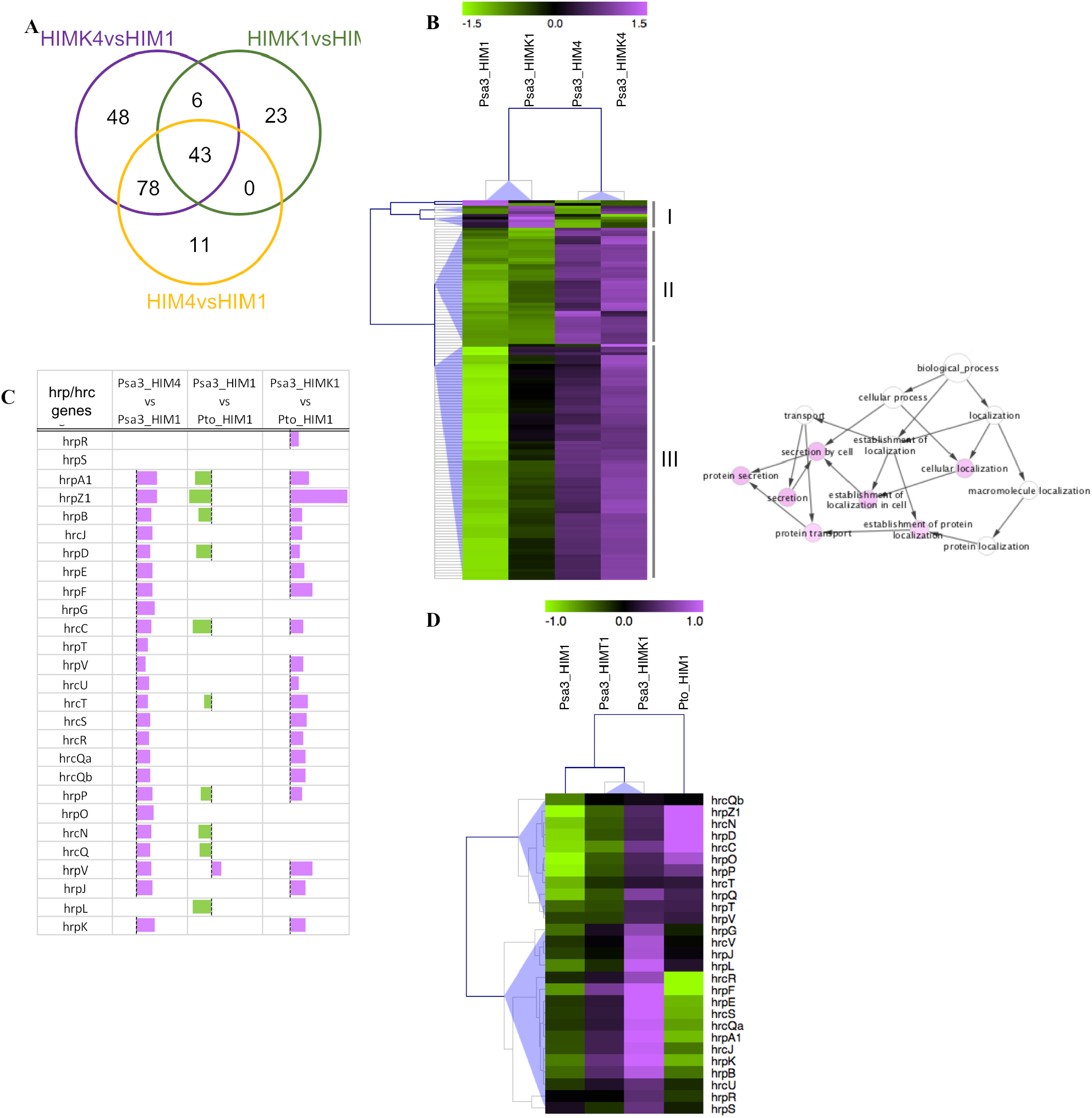
Kiwifruit extract anticipates T3SS-related gene expression and makes the induction of virulence in *P. syringae* pv. *actinidiae* (Psa) as fast as *P. syringae* pv. *tomato* (Pto) DC3000. (A) Common and specific genes upregulated in Psa3 incubated in minimal medium for 4 h (HIM4 *vs* HIM1) or in the presence of kiwifruit extract for 1 h (HIMK1 *vs* HIM1) or 4 h (HIMK4 *vs* HIM1). The Venn diagram was generated using Draw Venn Diagram. (B) Hierarchical clustering of the absolute expression level of Psa3 genes upregulated after 1 and 4 h in minimal (*hrp*-inducing) medium (HIM1 and HIM4) or HIM supplemented with kiwifruit extract (HIMK1 and HIMK4). Clusters were generated using MeV with normalized fluorescence values, and blue triangles indicate the different clusters. For each cluster (I–III), functional category enrichment was analyzed using BINGO (right panel). (C) Graphical representation of the differential expression of *hrp*/*hrc* genes in Psa3 grown in minimal medium for 1 or 4 h (Psa3_HIM1 or Psa3_HIM4) or in the presence of kiwifruit extract for 1 h (Psa3_HIMK1) and in Pto DC3000 grown in minimal medium for 1 h (Pto_HIM1). Each bar represents the log2 fold change of expression for the corresponding gene. (D) Hierarchical clustering of the absolute expression level of *hrp/hrc* genes in Psa3 and Pto DC3000 grown in minimal medium (Psa3_HIM1 and Pto_HIM1) and Psa3 grown in presence of kiwifruit (Psa3_HIMK1) or tomato (Psa3_HIMT1) extracts. Clusters were generated using MeV with normalized fluorescence values, and blue triangles indicate the different clusters.

The further comparison of T3SS-related gene expression between Psa3 and the model bacterium Pto DC3000 revealed that almost all *hrp/hrc* genes in Psa3 (24/27) were induced by incubation in minimal medium for 1–4 h, whereas none were induced in Pto DC3000 over the same period (**Supplementary Dataset S4**). Moreover, after 1 h in HIM, several *hrp/hrc* genes in Psa3 appeared to be downregulated compared to Pto DC3000 (**Fig. 2C**). This indicated that transcription levels were already elevated in Pto DC3000 after incubation for 1 h in HIM, whereas the same levels in Psa3 were only reached several hours later. This suggests the presence of a slower T3SS induction mechanism in Psa compared to the model strain Pto DC3000, which was supported by the hierarchical clustering of *hrp/hrc* genes showing that the transcript level of most genes (cluster I) was higher overall in Pto DC3000 (Pto_HIM1) than Psa3 (Psa3_HIM1), with only a few genes in cluster II showing different behavior (**Fig. 2D**). The faster response of Pto DC3000 to minimal medium was also supported by the very small number of genes modulated in this strain after incubation for 1 h in the presence of kiwifruit or tomato extracts, or incubation for 1–4 h in the apoplast-like medium (**Supplementary Datasets S5– S7**).

Interestingly, the expression level of *hrp/hrc* genes in Psa3 was similar to or higher than that in Pto DC3000 when grown in presence of the kiwifruit extract, with 20/27 genes showing a significantly higher expression level in Psa3. This supports the presence of kiwifruit-derived signals that trigger Psa3 to induce the rapid expression of T3SS-related genes, reaching or exceeding the transcript levels observed in Pto DC3000 but at an earlier stage (**Fig. 2C**). A similar trend was observed in the presence of tomato extract, although the genes were modulated to a lesser extent compared to the kiwifruit extract (**Fig. 2D**). Overall, these data show that T3SS induction is faster in Pto DC3000 than Psa3 under apoplast-like conditions, but that plant signals can anticipate T3SS activation in Psa3.

### Protein secretion by Psa3 is enhanced by plant extracts *in vitro*

To determine whether the fast response of Psa3 to plant signals correlates with T3SS functionality, we analyzed the profile of Psa3 proteins secreted *in vitro*. Initially, we tested the bacteria at 28 °C, the optimal growth temperature used for transcriptomic analysis in this study and previous reports (4). Surprisingly, although many genes related to protein secretion were induced in minimal medium at this temperature, very few proteins overall were detected in the extracellular medium, regardless of the presence or absence of kiwifruit extract (**Fig. 3A**). Interestingly, a greater number of secreted proteins were detected at 18 °C, close to the optimal infection temperature for Psa. Furthermore, in line with the upregulation of T3SS-related genes described above, the number of secreted proteins, and the overall signal intensity increased in the presence of kiwifruit extract. Similarly, the kiwifruit extract significantly increased the number of proteins secreted by Psa1 at 18 °C (**Supplementary Fig. S2**, left panel), in line with the greater *hrpA1* promoter activity reported in response to kiwifruit extract in this biovar (4). In contrast, the Psa2 secreted protein profile featured a larger number of secreted proteins at 28 °C and showed no response to the kiwifruit extract (**Supplementary Fig. S2**, right panel), in accordance with the different behavior of Psa2 reflecting its inability to induce the T3SS (4). Given that *hrpA1* promoter activity was not influenced by temperature (**Supplementary Fig. S3**), the observed temperature-dependent protein secretion may be related to secretion system assembly or functionality. Likewise, the addition of tomato extract significantly increased the total number of proteins secreted by Psa3 compared with cells incubated in minimal medium at 18 °C (**Fig. 3B**).

**Fig. 3.**
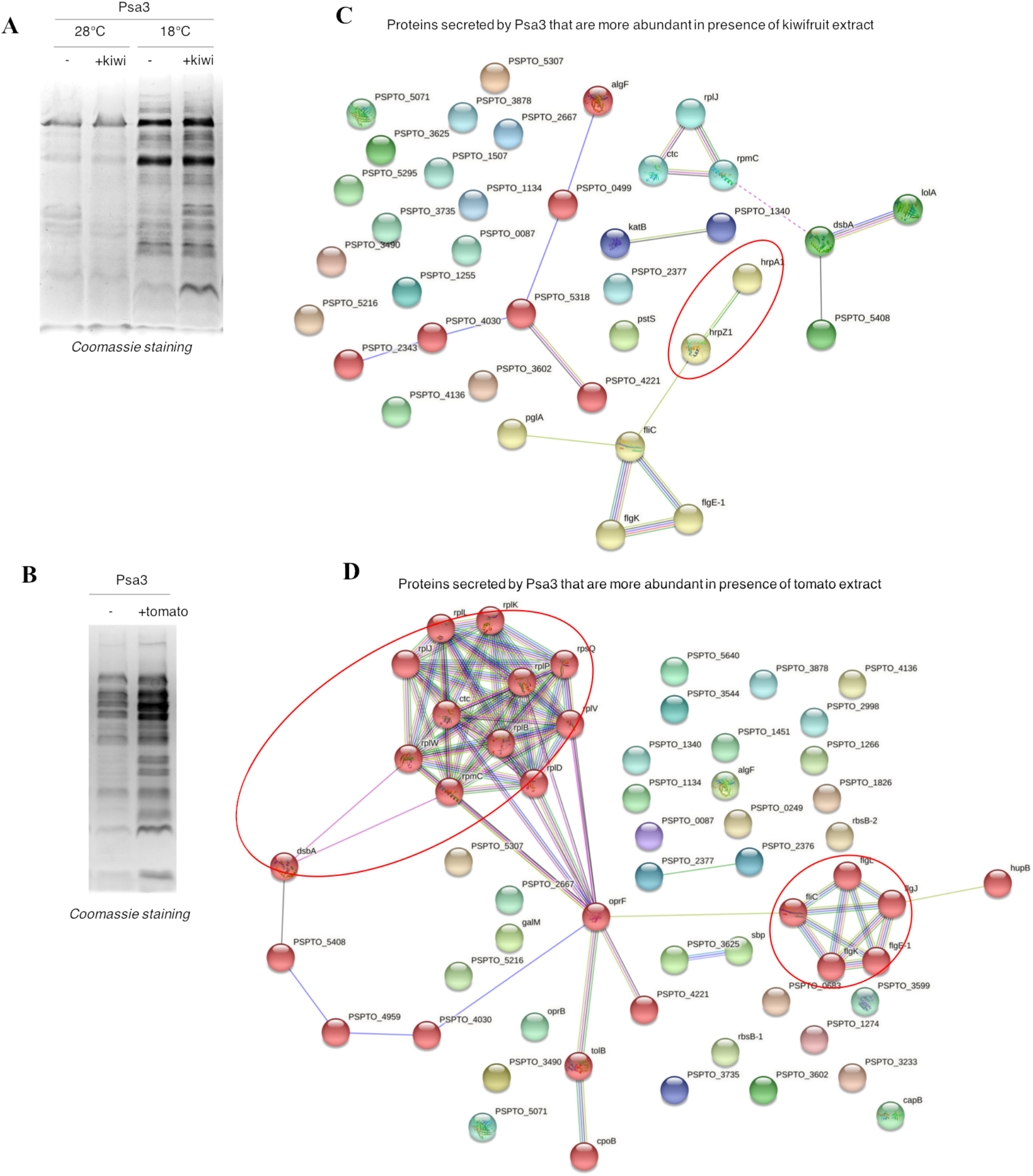
Kiwifruit and tomato extracts increase *P. syringae* pv. *actinidiae* (Psa) protein secretion *in vitro* in a temperature-dependent manner. (A) Comparison of the *in vitro* secretomes of Psa3 grown in minimal medium (HIM) with (+kiwi) or without (-) kiwifruit extract at 28 or 18 °C. (B) Comparison of the *in vitro* secretomes of Psa3 grown in minimal medium (HIM) with (+tomato) or without (-) tomato extract at 18°C. (C,D) Functional relationships among proteins secreted by Psa3 in the presence of (C) kiwifruit or (D) tomato extracts, analyzed using STRING.

We identified 195 secreted proteins that differed in abundance based on secretome analysis by mass spectrometry (**Supplementary Dataset S8-S9**). Semi-quantitative proteomic analysis showed that the abundance of 81 proteins was significantly modulated in the presence of kiwifruit extract (33 up and 48 down) and 114 proteins were similarly affected in the presence of tomato extract (56 up and 58 down) in each case compared to HIM, with 66 proteins affected by both extracts (**Supplementary Fig. S4**). Potential functional interactions among these proteins were investigated using STRING (**Fig. 3C,D**). Kiwifruit extract significantly increased the secretion of HrpA1 (FC = +6.47) and HrpZ1 (FC = +2.29), two of the three most abundant harpins secreted by the T3SS involved in effector transport (20). These proteins were included in a cluster with proteins involved in the flagellum-related T3SS and the flagellin protein FliC (**Fig. 3C, red cluster**). Interestingly, the latter were also secreted more abundantly in the presence of tomato extract (**Fig. 3D, green cluster**). Moreover, in line with the transcriptome profile, the tomato extract induced the secretion of proteins related to translation (yellow cluster), and proteins involved in the Pal-Tol system, which plays a role in the T3SS and flagellum-mediated virulence in enterohemorrhagic *Escherichia coli* (21). Notably, both extracts also induced the secretion of the disulfide bond-forming protein DsbA, which is required for the T3SS to function in Pto DC3000 (19). The kiwifruit extract did not modify the secreted protein profile in PtoDC3000 or Psa2, but the abundance of some secreted proteins in both strains was modulated by the tomato extract (**Supplementary Fig. S5**).

### Earlier T3SS induction by plant signals influences the triggering of hypersensitive cell death by Psa3 in *A. thaliana*

Psa3 is unable to induce an hypersensitive response in *A. thaliana* ecotype Col-0 (22) (**Supplementary Fig. S6A**), even though it produces the AvrRpm1 effector recognized by the host receptor RPM1. However, this does not reflect the inability of AvrRpm1Psa to trigger an RPM1-dependent response (23) because a transformed Psa3 strain carrying the well-known avirulence factor AvrB (Psa3-AvrB) still failed to induce hypersensitive cell death at 24 °C, the typical temperature used for ion leakage experiments, or at 18 °C, which is required for efficient protein secretion by Psa3 (**Supplementary Fig. S6B**). Interestingly, the pre-incubation of Psa3-AvrB in HIM for 1 h before infiltration in *A. thaliana* Col-0 led to the induction of a significant hypersensitive response, which was enhanced if kiwifruit extract was present in the pre-incubation medium (**Fig. 4A-B**). This demonstrates that induction of the T3SS under apoplast-like conditions and its enhancement by kiwifruit extract, at the mRNA and protein levels, leads to a functional T3SS that can efficiently translocate effectors into *A. thaliana* Col-0. Like protein secretion *in vitro*, the hypersensitive response induced by Psa3-AvrB is strictly temperature dependent, occurring only when infected leaf disks are incubated at 18 °C. In contrast, the pre-incubation temperature does not prevent Psa3-AvrB from triggering the hypersensitive response (**Fig. 4B**). Similarly, when Psa1 carrying the avirulence factor AvrB was pre-incubated for 1 h in HIM, a hypersensitive response was induced in *A. thaliana* Col-0, and was enhanced in the presence of kiwifruit extract (**Fig. 4C**). Remarkably, however, pre-incubation did not allow Psa2-AvrB to trigger a hypersensitive response, showing that hypersensitive cell death induced by Psa3 (and Psa1) is linked to the activation or enhancement of the T3SS, which does not occur in Psa2. Tomato extract also enhanced the hypersensitive response induced by Psa3 (**Fig. 4D**), agreeing with the induction of the T3SS at the mRNA level. However, although the treatment with tomato extract increases the number of secreted proteins also in Pto DC3000 and Psa2, it does not enhance the hypersensitive response induced by the same strains carrying AvrB (**Supplementary Fig. S5, S7**). This confirms that the enhanced hypersensitive response triggered by Psa3 is dependent on the ability of plant extracts to stimulate the T3SS.

**Fig. 4.**
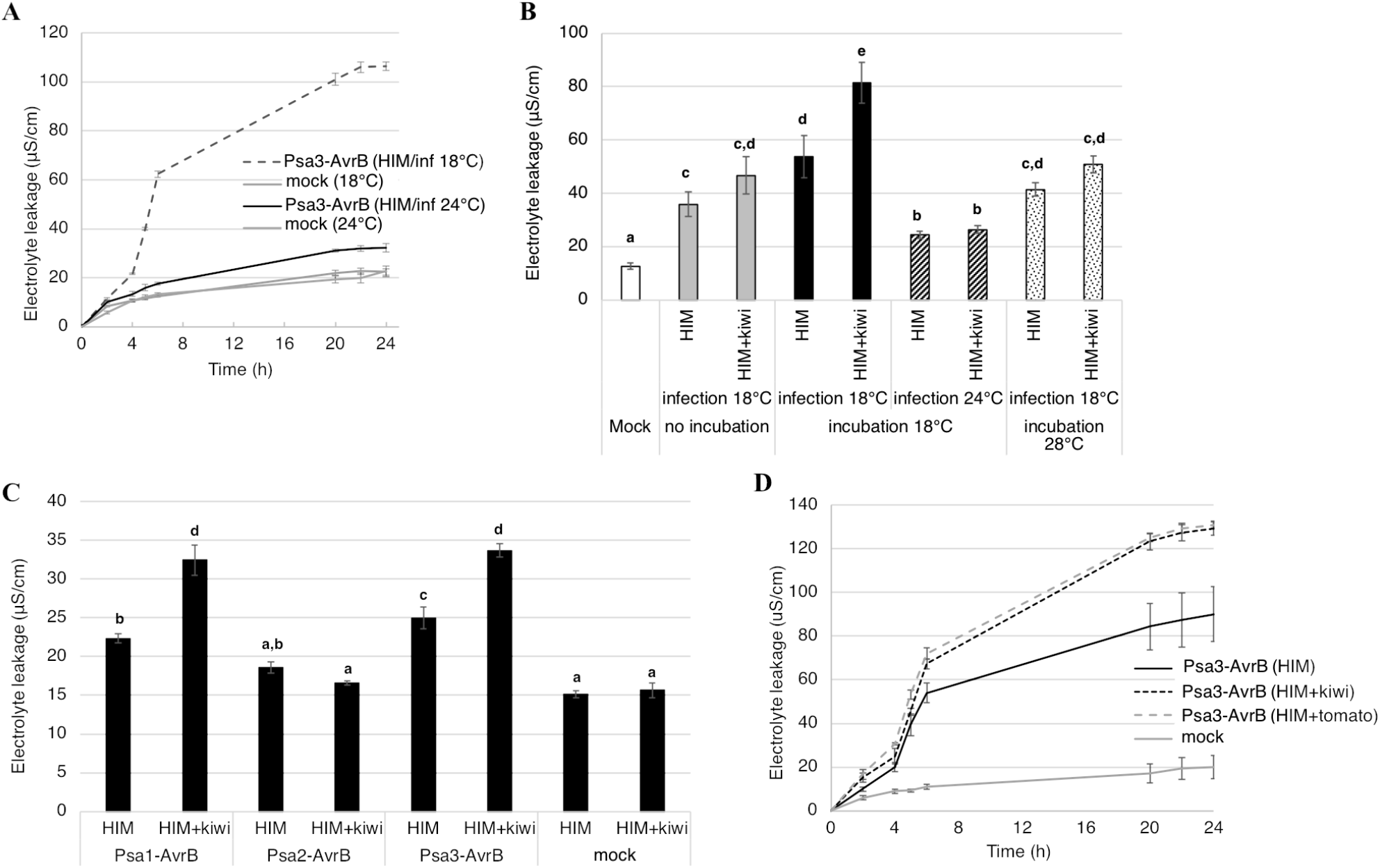
Psa3 pre-incubated under apoplast-like conditions with kiwifruit extract induces temperature-dependent hypersensitive cell death in the non-host plant *Arabidopsis thaliana* Col-0. (A) Time course of electrolyte leakage in *A. thaliana* Col-0 leaf disks infiltrated with *P. syringae* pv. *actinidiae* biovar 3 (Psa3) carrying the *AvrB* gene and pre-incubated in minimal medium (HIM) prior to infiltration. Infected leaf disks were incubated at 18 or 24 °C as indicated. Mock-infiltrated leaf disks were used as negative controls. Values correspond to the mean of three biological replicates, each including three technical replicates, ± standard error. (B,C) Electrolyte leakage 6 h postinfiltration in *A. thaliana* Col-0 leaf disks inoculated with (B) Psa3 carrying the *AvrB* gene preincubated (or not) for 1 h in HIM or in the presence of kiwifruit extract (HIM+kiwi), or (C) different Psa biovars (Psa1, Psa2 and Psa3) carrying the *AvrB* gene. In (B), the bacteria were pre-incubated at 18 or 28 °C and infected leaf disks were incubated at 18 or 24 °C, as indicated. In (C), Psa strains were pre-incubated for 1 h in HIM or in the presence of kiwifruit extract (HIM+kiwi) at 18 °C and infected leaf disks were incubated at 18°C. Values correspond to the mean of three biological replicates, each including three technical replicates, ± standard error. Different letters indicate statistically significant differences according to analysis of variance (ANOVA, p < 0.05). (D) Time course of electrolyte leakage in *A. thaliana* Col-0 leaf disks infiltrated with Psa3 carrying the *AvrB* gene and pre-incubated for 1 h in HIM, or HIM supplemented with kiwifruit (HIM+kiwi) or tomato (HIM+tomato) extracts prior to infiltration. The bacteria were pre-incubated at 18 °C and leaf disks were incubated at the same temperature. Mock-infiltrated leaf disks were used as negative controls. Values correspond to the mean of three biological replicates, each including three technical replicates, ± standard error. Different letters indicate statistically significant differences according to ANOVA (p < 0.05).

## Discussion

The role of host plant signals in the induction of Psa virulence is not completely understood, and the early phase of the bacterial response has not been studied in detail. We, therefore, profiled the transcriptome of Psa3 incubated in minimal medium with and without kiwifruit or tomato leaf extracts for 1 h. The presence of the kiwifruit extract led to the specific and strong induction of genes related to the T3SS, particularly the syringe structure (helper and secretion proteins), after only 1 h, suggesting that the host interaction anticipates T3SS stimulation. The tomato extract also induced *hrp* gene expression, but the response was less intense compared to the kiwifruit extract. Our results highlight the capacity of host plants to trigger the early activation of the T3SS. This indicates that the molecular analysis of T3SS induction may provide important information about the capacity of a bacterial strain to infect plants, even if a natural infection has yet to be reported.

Despite the presence of at least 30 effectors per strain and many pathovars, Psa still appears to have a limited host range. For example, the disease caused by Psa develops slowly in *A. thaliana* if at all (24). Moreover, effector-triggered immunity (ETI) in *A. thaliana* is pervasive following inoculation with Psa because 97% of Psa strains carry one or more immunity-eliciting effectors and 95% of Psa strains are recognized by the host due to two prominent R proteins (ZAR1 and CAR1). This demonstrates why the presence/absence of effectors and/or R proteins is necessary but not sufficient to predict the outcome of interactions between plants and bacteria. Indeed, Psa possesses the effector HrpZ5, which induces a hypersensitive response in *A. thaliana* Col-0 when introduced by Pto DC3000 but not by Psa, as shown herein and in previous studies (22). Moreover, the genetic transformation of Psa with the well-characterized avirulence gene *AvrB* does not significantly improve the induction of a hypersensitive response. Therefore it appears that Psa is likely unable to transfer its effectors efficiently, at least in *A. thaliana* Col-0. Interestingly, the pre-incubation of Psa in minimal medium allowed the induction of a hypersensitive response by the modified Psa3-AvrB strain, which was enhanced by adding kiwifruit extract to the medium, demonstrating that the enhancement of T3SS induction prior to the infiltration of *A. thaliana* Col-0 allows the assembly of a functional T3SS that can transfer effectors recognized by the corresponding R proteins and thus induce an efficient hypersensitive response.

As in a real dialogue, the interaction between pathogens and plants involves the mutual sensing of signals followed by appropriate responses. According to the current zig-zag model, plants first develop a general response based on pathogen-associated molecular patterns (PAMPs), known as PAMP-triggered immunity (PTI), to inhibit microbial colonization of the apoplast. Successful bacterial pathogens counter this response by using the T3SS to introduce PTI-suppressing type III effectors (25–27) that were acquired long before the advent of agriculture and even before the speciation of some of their host plants (28). However, PTI can directly or indirectly inhibit the introduction of type III effectors (29) and, when induced by flagellin in non-host plants, PTI can suppress the hypersensitive response elicited by *P. syringae* pv. *averrhoi* (30). Virulent bacteria must rapidly overcome T3SS restriction by PTI, allowing them to transfer their effectors and initiate effector-triggered susceptibility (ETS). In compatible interactions (which lead to the expression of disease symptoms), the initially low quantity of effectors transferred to the plant may be able to suppress PTI enough to allow the transfer of further effectors (29). However, this would mean that the failure of avirulent bacteria would rely on the absence of effectors that can block PTI, which is unrealistic considering the large effector repertoire of phytopathogenic bacteria. It is therefore reasonable to assume that the timing of bacterial virulence and plant defense responses may be another fundamental parameter governing the outcome of such interactions. In agreement with this hypothesis, three pathovars of *Pseudomonas savastanoi* (namely *fraxinii, savastanoi* and *nerii* showed different expression profiles for the four main T3SS-related genes when incubated in minimal medium, possibly accounting for the differences in their virulence (31). This may play a role in the ability of pathogenic strains to overcome PTI-mediated T3SS inhibition and inject virulence effectors that lead to the development of disease symptoms, as shown for the Pto DC3000/*A. thaliana* and *P. syringae* pv. *tabaci* (Pta)/tobacco interactions. Accordingly, we found that Pto DC3000 was able to fully activate its T3SS within 1 h.

As well as inducing T3SS-related gene expression, the plant extracts also appear to act at the protein level by stabilizing some of the T3SS components, such as HrpZ1. Indeed, the kiwifruit and tomato extracts both increased the abundance of DsbA without affecting *dsbA* mRNA levels. A similar correlation was observed in *A. thaliana* mutants with a modified salicylic acid (SA) pathway during interactions with PtoDC3000, resulting in higher levels of HrpZ1 and DsbA after 24 and 48 h, but no change in *dsbA* gene expression (19). Interestingly, a *P. syringae dsbA* mutant was less virulent and showed partial impairment of the T3SS (32). Protein secretion through the T3SS still occurs in this background, but the efficiency is lower, as also observed in a *dsbA* mutant strain of *Yersinia pestis* (33). The accumulation of DsbA in response to plant extracts may therefore contribute to the higher efficiency of protein secretion through the T3SS under the same conditions that induce T3SS-related gene expression.

We found evidence that temperature is a key factor governing the balance between virulence and plant defense. Two recent studies revealed transcriptomic and proteomic differences between virulent and avirulent forms of Pto DC3000 during interactions with *A. thaliana* (19). However, the experimental design did not consider the role of temperature in the outcome, regardless of the presence/absence of avirulence factors. Nevertheless, the role of temperature has been described and it clearly regulates the efficiency of defense responses in plants (34,35). Based on our hypothesis in which temperature is a key regulator of the T3SS, we used the Psa3-AvrB strain to monitor the induction of a hypersensitive response *A. thaliana* Col-0 and found that the temperature of infection was more important than the temperature of pre-incubation for T3SS induction, and thus T3SS functionality. In this pathosystem, programmed cell death is triggered at 18 °C suggesting that the Psa T3SS is more efficient than the PTI response (34), and our analysis of *in vitro* protein secretion provided supportive evidence. Interestingly, the hypersensitive response triggered by Pto DC3000 is not temperature dependent. However, high temperatures promote Pto DC3000 effector translocation, probably by inhibiting the SA-mediated pathway (36), in line with the greater abundance of HrpZ1, HrpK1, and HrpW in *A. thaliana sid2/pad4* mutants lacking this pathway (19). However, the more efficient translocation of type III effectors, which could contribute to a stronger hypersensitive response, would be balanced by the lower capacity of the plant to induce a hypersensitive response in the absence of SA-mediated signals (37). In the same manner, the production of the phytotoxin coronatine by Pto DC3000 is not temperature dependent, but in *P. syringae* pv. *glycinea* (Psg), it is induced only at low temperatures (38). Overall, this highlights the peculiar behavior of Pto DC3000, which differs at least from Psa3 and Psg by inducing virulence regardless of the temperature. In this context, it would be interesting to evaluate the dynamic T3SS activation process in different strains of the *Pseudomonas* complex to determine whether this may affect the outcome of the interactions with different plant species, depending on the capacity of plants to induce rapid defense responses that restrict the T3SS before its induction.

Our work demonstrates the need to consider not only the presence/absence of a bacterial effector repertoire combined with the presence/absence of the corresponding targets/genes in the host plant (leading to susceptibility and disease, respectively), but also the capacity (or opportunity) for plants to activate their basal defense mechanisms or for bacteria to express and assemble an efficient T3SS for the injection of virulence effectors. In this scenario, the aim of the arms race between plants and bacteria is not only to evolve new weapons (i.e., the acquisition of new genomic features) but also to gain the capacity to mobilize the existing ones more rapidly thus mounting an effective defense (plants) or enhancing virulence (bacteria). The temperature would play a key role in this competition, suggesting that the outcome of the molecular interaction with the plant is highly contingent on the biotic and abiotic context. Overall, our findings may explain, at least in part, the overlapping host range continuum for strains of the Psa complex (39).

## Materials and methods

### Bacterial strains and media

Single colonies of *Pseudomonas syringae* pv. *actinidiae* CRA-FRU 8.43 (biovar 3), J35 (biovar 1), KN.2 (biovar 2) and *Pseudomonas syringae* pv. *tomato* DC3000 (Pto) grown on rich solid medium (KB agar) were inoculated into KB medium and incubated overnight at 28 °C, shaking at 200 rpm. When cells reached the late log phase, they were collected by centrifugation (5000 × g, 10 min, room temperature), washed three times in liquid *hrp*-inducing medium (HIM) (8), and resuspended at a final optical density (OD600) of 0.1 in HIM with or without kiwifruit or tomato leaf extract at a final concentration of 1%. After incubation for 1 or 4 h at 28 °C, shaking at 200 rpm, the cells were harvested by centrifugation as above and the pellets (~2.4 × 10^9^ cells) were stored at −20°C.

### Preparation of leaf extracts

Leaves and petioles removed from 20 kiwifruit (*Actinidia deliciosa* ‘Hayward’) or tomato (*Solanum lycopersicum*) plants cultivated at 24 °C with 60% relative humidity were ground in a kitchen juice extractor and squeezed on ice. The collected raw extract was centrifuged repeatedly (5000 rpm, 10 min, 4 °C) until clarified, and the resulting supernatant was passed through a 0.2-μm sterilization filter before storage at –20 °C.

### RNA extraction and microarray chip hybridization

RNA was extracted from each sample using the Spectrum Plant Total RNA Kit (Sigma-Aldrich). Residual DNA was removed using the TURBO DNA-free kit (Thermo Fisher Scientific, Waltham, MA, USA). RNA concentrations were determined using a Nanodrop 2000 spectrophotometer (Thermo Fisher Scientific). First-strand cDNA was synthesized from 1 μg total RNA using the SuperScript III Reverse Transcriptase enzyme kit (Invitrogen, Carlsbad, CA, USA). Samples were labeled for microarray analysis using the One-Color Microarray-Based Gene Expression Analysis Low Input Quick Amp WT Labeling kit (Agilent Technologies, Santa Clara, CA, USA), according to the manufacturer’s instructions.

### Microarray and data analysis

The bacterial transcriptome was interrogated using a custom SurePrint G3 GE 8×60K chip designed in house and produced by Agilent Technologies (4). The fluorescence intensity for each probe was measured using an Agilent G4900DA SureScan Microarray Scanner System with Agilent Scan Control software, and data were extrapolated using Agilent Feature Extraction software. Fluorescence intensities were calculated by robust multi-array averaging, including adjustment for background intensity, log2 transformation, and quantile normalization. Normalized data were used to identify differentially expressed genes (DEGs) with threshold values of P < 0.05 and log2 FC values > |1|. DEGs were compared across different strains and/or conditions using the online software Calculate and Draw Custom Venn Diagrams (http://bioinformatics.psb.ugent.be/webtools/Venn/).

### Measurement of hypersensitive cell death in *A. thaliana*

Hypersensitive cell death was assessed by measuring electrolyte leakage from infected cells (12). Bacterial strains carrying the avirulence gene *AvrB* were grown on KB agar and single colonies were used to inoculate 30 ml of KB liquid medium supplemented with 50 μg/ml rifampicin (PtoAvrB only) and 50 μg/ml kanamycin (all strains). After overnight incubation at 28 °C, shaking at 200 rpm, the suspensions were centrifuged (5000 rpm, 15 min, room temperature) and the pellets were washed three times with 10 mM MgCl_2_. The bacteria were resuspended in 10 mM MgCl_2_, HIM or HIM+1%kiwi (final volume = 20 ml) at OD_600_ = 0.1. Three 8-mm leaf discs (representing three different leaves) from six *A. thaliana* Columbia-0 (Col-0) plants were vacuum infiltrated with the bacterial suspensions, or with 10 mM MgCl_2_, HIM or HIM+kiwi as negative controls. All infiltrated leaf discs were washed in milli-Q water for 30 min with gentle agitation (90 rpm) and distributed to three wells (six disks/well) containing 2 ml of milli-Q water. The three wells acted as three technical replicates for each condition. Conductivity (μS/cm) was measured after 0, 2, 4, 6 and 24 h. For each time point and each condition, conductivity was calculated as the average of the values of the three technical replicates and the values at time 0 were subtracted for normalization.

### Analysis of *hrpA1* promoter activity

Psa strain CRAFRU8.43 carrying the *phrpA1::gfp* construct (13) was grown on KB agar, and single colonies were used to inoculate 15 ml of KB liquid medium supplemented with 40 μg/ml gentamicin followed by overnight incubation at 28 °C, shaking at 200 rpm. The cells were recovered by centrifugation (5000 rpm, 15 min, room temperature) and were resuspended in fresh HIM with or without kiwifruit or tomato leaf extract (1% final concentration) at OD_600_ = 0.1. We transferred 200 μl of bacterial cells to the wells of transparent 96-multiwell plates and GFP fluorescence was measured every 15 min for 8 h using an Infinite 200 Pro fluorimeter (Tecan, Männedorf, Switzerland) at λ_exc_ = 485 nm and λ_em_ = 535 nm. The fluorescence value at time zero was subtracted from the values at other time points for normalization.

### Hierarchical clustering

Normalized fluorescence intensities from microarray experiments were imported as data matrices into MeV (14). The data were adjusted as median center genes/rows and clustered using the hierarchical clustering module. Gene and sample trees were clustered with optimized gene and sample leaf orders using Pearson correlation and average linkage clustering. The trees were subsequently cut into clusters using a distance threshold (0.5–1) empirically adjusted to highlight the most relevant features of the trees.

### Collection of secreted proteins and SDS-PAGE analysis

Bacterial strains were grown on KB agar and single colonies were used to inoculate 50 ml of KB liquid medium, followed by overnight incubation at 28 °C, shaking at 200 rpm. Cell suspensions were centrifuged (5000 rpm, 10 min, room temperature) and the pellets were washed three times in HIM before resuspension in 50 ml HIM with or without kiwifruit or tomato extract (final concentration = 1%) at a final OD_600_ of 0.8. Bacteria were incubated for 24 h at 18 or 28 °C, shaking at 200 rpm. Following another round of centrifugation as above, the supernatants were concentrated 50-fold using Amicon Ultra-4 centrifugal filters with a molecular weight cut-off of 3 kDa (Merck-Millipore, Burlington, MA, USA). Samples were mixed with 3× Laemmli loading buffer and heated for 3 min at 85 °C before SDS-PAGE analysis. Polyacrylamide gels were stained with Coomassie Brilliant Blue R-250 for protein visualization.

### Analysis of secreted proteins by LC-MS/MS

Supernatants obtained as described above were precipitated overnight at –20 °C with four volumes of ice-cold acetone. After centrifugation (14,000 rpm, 10 min, 4 °C), the pellets were air-dried for 10 min and resuspended in solubilization solution comprising 7 M urea, 2 M thiourea, 3% (w/v) CHAPS, 40 mM Tris, and Complete Mini protease inhibitor cocktail (Roche, Basel, Switzerland). Proteins were quantified using the Bradford method (15). Before LC-MS/MS analysis, secreted proteins were digested with trypsin (Applied Biosystems, Waltham, MA, USA) as previously described (16). Comparative proteomic analysis by LC-MS/MS was carried out using a micro-LC device (Eksigent Technologies, Dublin, OH, USA) coupled to a 5600+ TripleTOF mass spectrometer (AB Sciex, Concord, ON, Canada). After peptide separation on a Halo Fused C18 column (AB Sciex) using a 30 min gradient of acetonitrile in 0.1% formic acid, MS and MS/MS spectra were collected as previously described (17). Briefly, data-dependent analysis (DDA) was combined with cyclic data-independent analysis (DIA) using a 25-Da window (36 sweeps in total) for identification and label-free quantification. Following two DDA and three DIA acquisitions with Analyst TF v1.7 (AB Sciex), the DDA files were searched using Protein Pilot v4.2 (AB Sciex) and Mascot v2.4 (Matrix Science, Boston, MA, USA). Mascot was set to allow two missed cleavages, the instrument was set to ESI-QUAD-TOF mode, carbamidomethyl cysteine was specified as a fixed modification, and oxidized methionine as a variable modification. Triplicate SWATH-MS scans were carried out for each sample, and the data were imported into Skyline for label-free quantification and the identification of secreted proteins. The peptide mass tolerance was set to 0.08 Da and the MS/MS tolerance to 10 ppm. The search was set to a monoisotopic mass with charges of 2^+^, 3^+^ and 4^+^. We searched a UniProt and Swiss-Prot reviewed database containing Psa group proteins as well as a target-decoy database, with the false discovery rate set to 1%.

### Protein–protein interactions analysis

Protein–protein interactions were predicted using the STRING database (http://string-db.org), including text-mining (18). The species under investigation was set to a medium confidence level (0.400) and we retrieved known and predicted interactions. Additional white nodes and network depth were kept to the minimum value of 1 to exclude as many false positive interactions as possible. Clusters were retrieved using MCL clustering with the inflation parameter set to 1.1.

## Acknowledgments

We acknowledge Dr. Marco Scortichini (Italian Council for Agricultural Research and Economics, Unit for Fruit Crop Research, CREA-FRC, Caserta, Italy) for providing the Psa strains CRAFRU 8.43, J35 and KN.2, and the Veneto Region for funding in the frame of the project “Progetto di innovazione per la difesa della pianta del kiwi e per la valorizzazione dei suoi frutti.” LR No. 32 del 9/08/1999 – art. 4 “Ricerca di interesse regionale e sperimentazione” (CUP) No. H16D14000090002.

## Author contribution

EV and AP conceived the work

MRP, DD, CC, JR, MM, EM performed the experiments

EV, DC, CS and FS analyzed and interpreted the data

EV wrote the original draft

All the authors revised and edited the manuscript

## References

1. O’Brien HE, Thakur S, Guttman DS. Evolution of Plant Pathogenesis in Pseudomonas syringae: A Genomics Perspective. Annu Rev Phytopathol. 2011 Sep 8;49(1):269–89.

2. Lindeberg M, Cunnac S, Collmer A. Pseudomonas syringae type III effector repertoires: last words in endless arguments. Trends Microbiol. 2012 Apr 1;20(4):199–208.

3. Hauser AR. The type III secretion system of Pseudomonas aeruginosa: infection by injection. Nat Rev Microbiol. 2009;7(9):654–65.

4. Vandelle E, Colombo T, Regaiolo A, Maurizio V, Libardi T, Puttilli M-R, et al. Transcriptional Profiling of Three Pseudomonas syringae pv. actinidiae Biovars Reveals Different Responses to Apoplast-Like Conditions Related to Strain Virulence on the Host. Mol Plant-Microbe Interactions®. 2021 Apr 1;34(4):376–96.

5. Galán JE, Collmer A. Type III Secretion Machines: Bacterial Devices for Protein Delivery into Host Cells. Science. 1999 May 21;284(5418):1322.

6. Xie Y, Shao X, Deng X. Regulation of type III secretion system in Pseudomonas syringae. Environ Microbiol. 2019 Dec 1;21 (12):4465–77.

7. Rahme LG, Mindrinos MN, Panopoulos NJ. Plant and environmental sensory signals control the expression of hrp genes in Pseudomonas syringae pv. phaseolicola. J Bacteriol. 1992;174(11):3499–507.

8. Rico A, Preston GM. Pseudomonas syringae pv. tomato DC3000 uses constitutive and apoplast-induced nutrient assimilation pathways to catabolize nutrients that are abundant in the tomato apoplast. Mol Plant-Microbe Interact MPMI. 2008;21(2):269–82.

9. Aldon D, Brito B, Boucher C, Genin S. A bacterial sensor of plant cell contact controls the transcriptional induction of Ralstonia solanacearum pathogenicity genes. EMBO J. 2000;19(10):2304–14.

10. Hernández-Morales A, De la Torre-Zavala S, Ibarra-Laclette E, Hernández-Flores JL, Jofre-Garfias AE, Martínez-Antonio A, et al. Transcriptional profile of Pseudomonas syringae pv. phaseolicola NPS3121 in response to tissue extracts from a susceptible Phaseolus vulgaris L. cultivar. BMC Microbiol. 2009;9:257–257.

11. Yu X, Lund SP, Scott RA, Greenwald JW, Records AH, Nettleton D, et al. Transcriptional responses of Pseudomonas syringae to growth in epiphytic versus apoplastic leaf sites. Proc Natl Acad Sci U S A. 2013;110(5):E425–34.

12. Imanifard Z, Vandelle E, Bellin D. Measurement of Hypersensitive Cell Death Triggered by Avirulent Bacterial Pathogens in Arabidopsis. In: De Gara L, Locato V, editors. Plant Programmed Cell Death: Methods and Protocols. New York, NY: Springer New York; 2018. p. 39–50.

13. Vandelle E, Puttilli MR, Chini A, Devescovi G, Venturi V, Polverari A. Application of Chemical Genomics to Plant–Bacteria Communication: A High-Throughput System to Identify Novel Molecules Modulating the Induction of Bacterial Virulence Genes by Plant Signals. In: Busch W, editor. Plant Genomics: Methods and Protocols. New York, NY: Springer New York; 2017. p. 297–314.

14. Howe EA, Sinha R, Schlauch D, Quackenbush J. RNA-Seq analysis in MeV. Bioinforma Oxf Engl. 2011/10/05 ed. 2011 Nov 15;27(22):3209–10.

15. Bradford MM. A rapid and sensitive method for the quantitation of microgram quantities of protein utilizing the principle of protein-dye binding. Anal Biochem. 1976 May 7;72:248–54.

16. Brandi J, Cheri S, Manfredi M, Di Carlo C, Vita Vanella V, Federici F, et al. Exploring the wound healing, anti-inflammatory, anti-pathogenic and proteomic effects of lactic acid bacteria on keratinocytes. Sci Rep. 2020 Jul 14;10(1):11572.

17. Dalla Pozza E, Manfredi M, Brandi J, Buzzi A, Conte E, Pacchiana R, et al. Trichostatin A alters cytoskeleton and energy metabolism of pancreatic adenocarcinoma cells: An in depth proteomic study. J Cell Biochem. 2018 Mar;119(3):2696–707.

18. Szklarczyk D, Gable AL, Lyon D, Junge A, Wyder S, Huerta-Cepas J, et al. STRING v11: protein-protein association networks with increased coverage, supporting functional discovery in genome-wide experimental datasets. Nucleic Acids Res. 2019 Jan 8;47(D1):D607–13.

19. Nobori T, Wang Y, Wu J, Stolze SC, Tsuda Y, Finkemeier I, et al. Multidimensional gene regulatory landscape of a bacterial pathogen in plants. Nat Plants. 2020 Jul 1;6(7):883–96.

20. Collmer Alan, Badel Jorge L., Charkowski Amy O., Deng Wen-Ling, Fouts Derrick E., Ramos Adela R., et al. Pseudomonas syringae Hrp type III secretion system and effector proteins. Proc Natl Acad Sci. 2000 Aug 1;97(16):8770–7.

21. Hirakawa H, Suzue K, Takita A, Awazu C, Kurushima J, Tomita H. Roles of the Tol-Pal system in the Type III secretion system and flagella-mediated virulence in enterohemorrhagic Escherichia coli. Sci Rep. 2020 Sep 23;10(1):15173.

22. Jayaraman J, Choi S, Prokchorchik M, Choi DS, Spiandore A, Rikkerink EH, et al. A bacterial acetyltransferase triggers immunity in Arabidopsis thaliana independent of hypersensitive response. Sci Rep. 2017 Jun 15;7(1):3557.

23. Yoon M, Rikkerink EH. *Rpa1* mediates an immune response to *avrRpm1Psa* and confers resistance against *Pseudomonas syringae* pv. *actinidiae*. Plant J. 2019;

24. Dillon MM, Almeida RND, Laflamme B, Martel A, Weir BS, Desveaux D, et al. Molecular Evolution of Pseudomonas syringae Type III Secreted Effector Proteins. Front Plant Sci. 2019 Apr;10(418).

25. Espinosa A, Alfano JR. Disabling surveillance: bacterial type III secretion system effectors that suppress innate immunity. Cell Microbiol. 2004 Nov 1;6(11):1027–40.

26. Chisholm ST, Coaker G, Day B, Staskawicz BJ. Host-Microbe Interactions: Shaping the Evolution of the Plant Immune Response. Cell. 2006 Feb 24;124(4):803–14.

27. Jones JDG, Dangl JL. The plant immune system. Nature. 2006 Nov 1;444(7117):323–9.

28. Morris CE, Moury B. Revisiting the Concept of Host Range of Plant Pathogens. Annu Rev Phytopathol. 2019 Aug 25;57(1):63–90.

29. Crabill E, Joe A, Block A, van Rooyen JM, Alfano JR. Plant Immunity Directly or Indirectly Restricts the Injection of Type III Effectors by the <em>Pseudomonas syringae</em> Type III Secretion System. Plant Physiol. 2010;154(1):233.

30. Wei C-F, Hsu S-T, Deng W-L, Wen Y-D, Huang H-C. Plant Innate Immunity Induced by Flagellin Suppresses the Hypersensitive Response in Non-Host Plants Elicited by Pseudomonas syringae pv. averrhoi. Yang C-H, editor. PLoS ONE. 2012 Jul 23;7(7):e41056.

31. Tegli S, Gori A, Cerboneschi M, Cipriani MG, Sisto A. Type Three Secretion System in Pseudomonas savastanoi Pathovars: Does Timing Matter? Genes. 2011;2(4):957–79.

32. Kloek AP, Brooks DM, Kunkel BN. A dsbA mutant of Pseudomonas syringae exhibits reduced virulence and partial impairment of type III secretion. Mol Plant Pathol. 2000 Mar 1;1(2):139–50.

33. Jackson MW, Plano GV. DsbA is required for stable expression of outer membrane protein YscC and for efficient Yop secretion in Yersinia pestis. J Bacteriol. 1999 Aug;181(16):5126–30.

34. Cheng C, Gao X, Feng B, Sheen J, Shan L, He P. Plant immune response to pathogens differs with changing temperatures. Nat Commun. 2013;4:2530–2530.

35. Menna A, Nguyen D, Guttman DS, Desveaux D. Elevated Temperature Differentially Influences Effector-Triggered Immunity Outputs in Arabidopsis. Front Plant Sci. 2015;6:995–995.

36. Huot B, Castroverde CDM, Velásquez AC, Hubbard E, Pulman JA, Yao J, et al. Dual impact of elevated temperature on plant defence and bacterial virulence in Arabidopsis. Nat Commun. 2017 Nov 27;8(1):1808.

37. Raffaele S, Rivas S, Roby D. An essential role for salicylic acid in AtMYB30-mediated control of the hypersensitive cell death program in Arabidopsis. FEBS Lett. 2006 Jun 12;580(14):3498–504.

38. Ullrich MS, Schergaut M, Boch J, Ullrich B. Temperature-responsive genetic loci in the plant pathogen Pseudomonas syringae pv. glycinea The GenBank accession numbers for the nucleotide sequences of mutants 560, 561, 562, 563, 564, 568, 570, 574, 590, 591, 593, 596, 599, 601, 605, 608, 613, 617, 618, 626 and 632 determined in this work are AF274322–AF274342, respectively. Microbiology. 2000 Oct 1;146(10):2457–68.

39. Morris CE, Lamichhane JR, Nikolić I, Stanković S, Moury B. The overlapping continuum of host range among strains in the Pseudomonas syringae complex. Phytopathol Res. 2019 Jan 16;1(1):4.

